# SvABA: Genome-wide detection of structural variants and indels by local assembly

**DOI:** 10.1101/105080

**Authors:** Jeremiah Wala, Pratiti Bandopadhayay, Noah Greenwald, Ryan O’Rourke, Ted Sharpe, Chip Stewart, Steve Schumacher, Yilong Li, Joachim Weischenfeldt, Xiaotong Yao, Chad Nusbaum, Peter Campbell, Matthew Meyerson, Cheng-Zhong Zhang, Marcin Imielinski, Rameen Beroukhim

**Author notes:** **Corresponding author**: Rameen Beroukhim, MD, PhD; Department of Cancer Biology, Dana-Farber Cancer Institute; 450 Brookline Avenue, Smith 1022, Boston, MA, 02115. Co-senior authors.

## Abstract

Structural variants (SVs), including small insertion and deletion variants (indels), are challenging to detect through standard alignment-based variant calling methods. Sequence assembly offers a powerful approach to identifying SVs, but is difficult to apply at-scale genome-wide for SV detection due to its computational complexity and the difficulty of extracting SVs from assembly contigs. We describe SvABA, an efficient and accurate method for detecting SVs from short-read sequencing data using genome-wide local assembly with low memory and computing requirements. We evaluated SvABA’s performance on the NA12878 human genome and in simulated and real cancer genomes. SvABA demonstrates superior sensitivity and specificity across a large spectrum of SVs, and substantially improved detection performance for variants in the 20-300 bp range, compared with existing methods. SvABA also identifies complex somatic rearrangements with chains of short (< 1,000 bp) templated-sequence insertions copied from distant genomic regions. We applied SvABA to 344 cancer genomes from 11 cancer types, and found that templated-sequence insertions occur in ~4% of all somatic rearrangements. Finally, we demonstrate that SvABA can identify sites of viral integration and cancer driver alterations containing medium-sized SVs.

## Introduction

Structural variants (SVs) are a broad class of genomic alterations that includes deletions, duplications and insertions, sequence inversions, and inter-chromosomal translocations, among other more complex topologies. SVs are an important source of variation in the human population (Sudmant et al. 2015) and create a significant amount of genomic disruption in cancer (Beroukhim et al. 2010;; Garraway and Lander 2013). In the germline, small to medium sized events between 10 bp - 10 Kbp are the primary source of structural variation (Mullaney et al. 2010). Large events (>10 Kbp) and inter-chromosomal translocations are rare in the germline due to stringent selection against gene dosage imbalance, but are prevalent in cancer, where genomes are often unstable and suffer frequent complex events(Stephens et al. 2009). The junctions connecting two SV breakpoints may also involve the insertion of novel bases created during DNA repair (Mahaney et al. 2009) or insertion of short fragments copied from elsewhere in the genome (Zhang et al. 2015; Liu et al. 2011).

Though inference from short-read alignments forms the core of most variant calling pipelines, properly aligning reads to SVs is particularly challenging due the substantial heterogeneity of SV sizes and topologies. In contrast to single-nucleotide variants (SNV) which affect only single base pairs (bps), SVs frequently involve long stretches of the genome and are often supported by reads that are completely or partially unaligned (soft-clipped). Alignment is particularly inaccurate at complex junctions that may contain sequences derived from more than two genomic loci, and at sites of integration of viral sequences, where the viral-supporting reads will be left completely unaligned. Finally, genomic repeats obscure the locations of true breakpoints, and can lead to false breakpoint calls, especially in the context of sequencing artifacts and homopolymer runs(Ross et al. 2013).

This diversity in SVs and indels has prompted the development of a number of alignment-based approaches and tools aimed at their detection. Small indels can often be inferred directly from the output of gapped read aligners as in BWA (Li 2013) or from local realignment of candidate reads as in Strelka (Saunders et al. 2012), GATK UnifiedGenotyper (DePristo et al. 2011), and FreeBayes (Garrison and Marth 2012). Longer indel variants can be obtained from direct realignment of clipped and unmapped reads, as in Pindel (Ye et al. 2009), or from targeted sequence assemblies such as in SOAPindel (Li et al. 2013), Scalpel (Narzisi et al. 2014), ScanIndel (Yang et al. 2015), BreaKmer (Abo et al. 2015) and laSV (Zhuang and Weng 2015). For larger SVs, the cornerstone of most alignment-based detection algorithms is the clustering of discordant mates, which are read pairs with insert-sizes and relative orientations that differ substantially from expectation based on the physical library preparation (Tuzun et al. 2005; Korbel et al. 2007). Discordant mate clustering is often followed by a more focused step to realign clipped reads at the candidate variant sites, such with DELLY (Rausch et al. 2012), Breakpointer (Drier et al. 2013), and Meerkat (Yang et al. 2013). Some tools like LUMPY perform inference directly from multi-part alignments(Layer et al. 2014).

Assembly-based approaches provide a fundamentally different approach to variant calling, with global whole-genome *de novo* assembly being the most comprehensive implementation. In principle, longer contigs assembled from short sequence reads can be more accurately aligned to the genome, enabling more sensitive detection of junction-spanning sequences supporting complex indels and SVs. However, whole-genome *de novo* assembly can be untenable in practice due to the large computational requirements, including significant amounts of memory (> 60 Gb for a 60x human genome) and substantial (> 1,000 hours) CPU time (Simpson and Durbin 2012). Additionally, while a number of variant detection tools have been designed using the alignment inference paradigm, SV detection from *de novo* assemblies is considerably less well studied.

In contrast to whole-genome global assembly, local assembly can be used to assemble only reads with an initial alignment to some loci in the reference, significantly reducing computational requirements. These have been applied in a targeted fashion to detect SVs and indels in exons (Scalpel (Narzisi et al. 2014)) and at sites of candidate SVs identified by alignment-based methods (TIGRA (Chen et al. 2014)). Local assemblies can also be applied genome-wide by assembling continuous small windows tiled across the entire genome. Genome-wide variant detection from local assemblies has been described for indel and SNP detection with Platypus (Rimmer et al. 2014) and HaplotypeCaller (DePristo et al. 2011), and has recently been described for SVs with NovoBreak (Chong et al. 2016).

Here we describe SvABA (structural variation analysis by assembly; “sah-bah”), a unified tool to efficiently detect SVs and indels genome-wide using local assembly. The basic idea of this approach is to perform local assembly to create consensus contigs from sequence reads with divergence from the reference, and to apply this procedure to every region of the genome. The contigs are then compared to the reference to annotate the variants. By uniting the different classes of variant-supporting reads into a single framework, we further expect that this assembly-first approach would be effective for variants of all sizes and require few parameters; in particular, SvABA can detect variants of sizes between small indels and large SVs, a gap not well covered by current SV analysis methods. In contrast to many SV and indel detection tools, SvABA depends only weakly on the quality of the initial alignments, and effectively realigns sequence reads to the reference with the aid of local sequence assembly. We further demonstrate SvABA’s ability to detect medium sized SVs and multi-part complex rearrangements, often using previously unmapped or poorly mapped sequence reads.

## Results

### Detection of SVs and indels with SvABA

SvABA assembles sequences from multiple classes of read alignments to discover SVs, indels, complex rearrangement junctions and sites of viral integration (Fig. 1a). Assembly is applied to small (e.g. 25 Kbp) overlapping local assembly windows covering the entire genome, or to a list of target regions (Fig 1b). In the read retrieval phase, SvABA extracts all sequence reads with significant divergence from the reference, including soft-clipped, gapped, discordant, and highly mismatched alignments, as well as unmapped sequences. These sequences are trimmed to remove low optical-quality bases and low-complexity sequences, and down-sampled at sites of high-coverage pileups (**Supplementary Fig. 1**). Candidate discordant reads are also realigned to the reference with BWA-MEM (Li 2013) to reduce false-positive discordant alignments. SvABA then uses discordant read clusters to connect distant local assembly windows together, which allows variant-supporting reads to be retrieved regardless of which end of the breakpoint they align to and increases the power to detect variants (see Methods).

**Figure 1:**
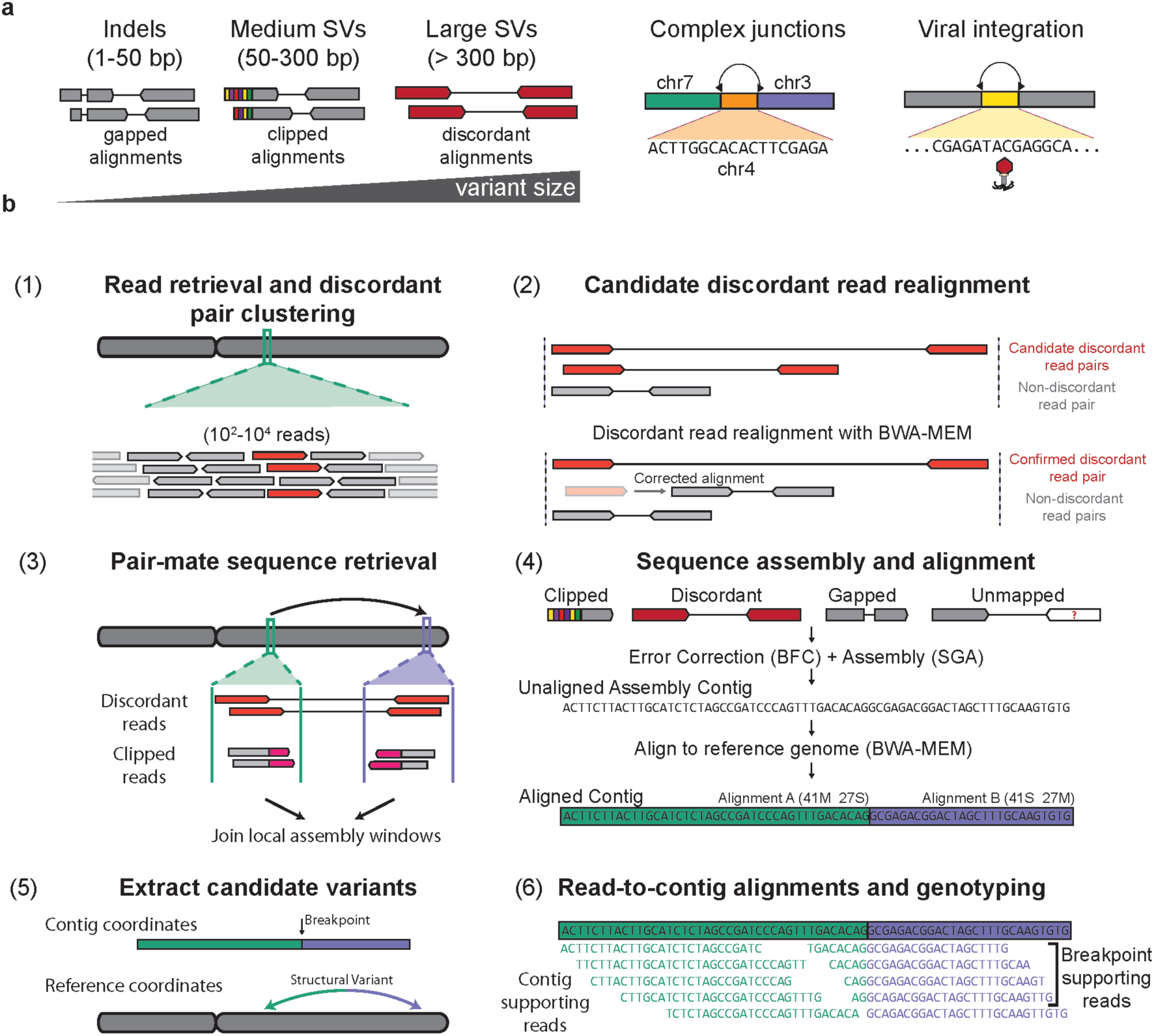
Overview of the SvABA structural variation detection tool. **a)** (left) SvABA assembles with SGA aberrantly aligned sequence reads that may reflect an indel or SV. Such reads include gapped alignments (for indels), clipped alignments (for medium and large SVs) and discordant read pairs (for large SVs). In additional to detecting indels and SVs, SvABA can identify complex rearrangement junctions (middle) and sites of viral integration (right). **b)** The workflow for the SvABA pipeline. (1) Reads within a small window are extracted from one or multiple BAM files and discordant reads are clustered. (2) Discordant reads are re-aligned to the reference to remove pairs that have a candidate non-discordant alignment. (3) The discordant read clusters are used to identify additional regions where reads should be extracted. (4) The sequences are error corrected with BFC and assembled with SGA into contigs. Contigs are immediately aligned to the reference with BWA-MEM. (5) Contigs with multi-part alignments or gapped alignments are parsed to extract candidate variants. (6) Sequence reads are aligned to the contig and to the reference to establish read support for the reference and alternative haplotypes.

Following read extraction and read clustering, the sequences are error corrected using BFC (Li 2015) and FM-indexed and assembled with String Graph Assembler (SGA) (Simpson and Durbin 2012). The contigs are then aligned to the reference genome using an in-memory integrated implementation of BWA-MEM (Li 2013). Contigs that align to the genome with gapped alignments produce candidate indels, and contigs with multi-part alignments produce candidate SVs (**Supplementary Fig. 2**). The level of support for candidate variants is obtained by aligning the raw read sequences to both the contig and reference genome using BWA-MEM. Variants are then scored based on the number of supporting reads and the quality of the contig alignment to the reference genome (see Methods).

SvABA can perform genome-wide local assembly and SV calling on a 30x genome with ~7 Gb of memory and ~40 CPU hours, orders of magnitude faster than global assemblies. The speed and efficiency of SvABA draws from the fusing and refactoring of several well-established C and C++ tools (SGA, htslib, BWA-MEM) into a single unified process, enabling in-memory manipulation of objects representing sequences, alignments, and assemblies with a minimal RAM and I/O footprint (Wala and Beroukhim 2016). SvABA can process local assembly windows in parallel, enabling a linear increase in speed through the use of additional CPU cores (set with a simple flag). The minimum input to SvABA consists only of a target reference genome and one or more alignment files (BAM, SAM or CRAM format) produced by standard alignment algorithms. The primary outputs are variant call files (VCF) representing the indels and SVs.

### Benchmarking SV calls with SVlib

Variant caller benchmarking remains a fundamental challenge in all areas of genomics (Zook et al. 2014; Parikh et al. 2016). Diversity in both the size and topology of variants presents a specific challenge for evaluation of SV calling performance. The ideal benchmarking pipeline would efficiently evaluate detection performance across this wide variant spectrum while remaining independent of the technical nuances of short-read sequencing and data processing. Simulated data have been previously used to evaluate detection performance for SVs(Layer et al. 2014), and offer a truth set for benchmarking caller sensitivity and specificity. However, current SV simulation approaches are typically limited to only a few classes and sizes of variants, and may not recapitulate the types of errors present in real sequencing data.

Alternatively, benchmarking on real data requires obtaining a validation callset based on orthogonal genotyping, sequencing and/or informatics methods. Examples of such validation approaches include PCR-based target enrichment followed by electrophoresis-based genotyping or Sanger sequencing, or long– or linked-sequencing. In particular, long-sequences such as obtained from whole genome *de novo* assembly or Pacific Biosciences (PacBio) sequencing, provide a promising and scalable truth set for validation. However, as with any method, the detection of SVs using these approaches is never free from false positives or false negatives, making it difficult for any to serve as an undisputed gold standard for validation.

With these considerations in mind, we developed SVlib, an SV benchmarking toolkit designed to address a wide range of variant sizes and classes. SVlib contains two primary modules. First, given a long-sequence alignment to a reference (e.g. long-read sequencing or *denovo* assembly contigs), SVlib will extract indel and SV variant calls and output to standard VCF format. SVlib can also genotype the extracted variants by efficiently aligning and scoring reads to the long-sequences. Second, SVlib provides the capability to simulate SVs ranging from single base pair indels to translocations, allowing for the creation of *in silico* genomes for benchmarking detection approaches. We grouped our analysis and benchmarking into three classes of variants (Fig. 1a), divided roughly by the types of evidence used for detection: small indels ≤ 50 bp, medium-sized deletions and duplications between 51 bp and 300 bp, and all SVs greater than 300 bp. We further assessed the ability of SvABA to correctly classify variants in cancer genomes.

### Sensitive detection of indels and SVs in NA12878

We used the widely studied NA12878 human genome to benchmark variant detection from a single germline sample. We ran SvABA on NA12878 whole-genome data sequenced to a mean coverage of 78.6-fold with 151 bp Illumina paired-end reads. Among variant-supporting contigs, the median contig length was 307 bp for indels (N50 = 330 bp) and 457 bp (N50 = 650) for SVs (**Supplementary Fig. 3**). SvABA identified 4,654 deletions > 50 bp, 1,894 duplications > 50 bp, 419 inversions, 149,117 small insertions (≤ 50 bp) and 178,632 small deletions (**Supplementary Data 1**). Among small variants (≤ 50 bp), 97.2% were represented in the dbSNP database. Agreement with dbSNP was highly size dependent, with 98.3% of variants ≤ 20 bp represented in the dbSNP database, compared with only 75.6% for variants between 20 - 50 bp (**Supplementary Fig. 4**). Despite this lower representation, 20 - 50 bp variants exhibited nearly the same level of read support (mean: 36.6 bp) as small variants (≤ 20 bp; mean: 39.5 bp), suggesting that true large indels are under reported in dbSNP.

SvABA integrated the different read signals to detect SVs using assembled contig realignment, discordant read clusters, or a combination of both signals. Detection of SVs from multi-part alignments of assembled contigs (as opposed to from discordant read alignments or gapped alignments) was a particularly important source of evidence for smaller variants (Fig. 2a). Among the SVs below 100 bp, 44.1% were identified through assembly-only, 2.9% from discordant reads only, and 52.9% having both signals. We thus find that a substantial amount of small structural variation can be found even in the absence of initial alignment-based signals. In contrast, 95.4% of larger SVs (> 300 bp) were support by discordant read clusters, with only 4.6% identified by assembly-only. Of these discordant-supported SVs, 86.5% were independently supported by realignment of assembly contigs, further boosting confidence for the variants.

**Figure 2:**
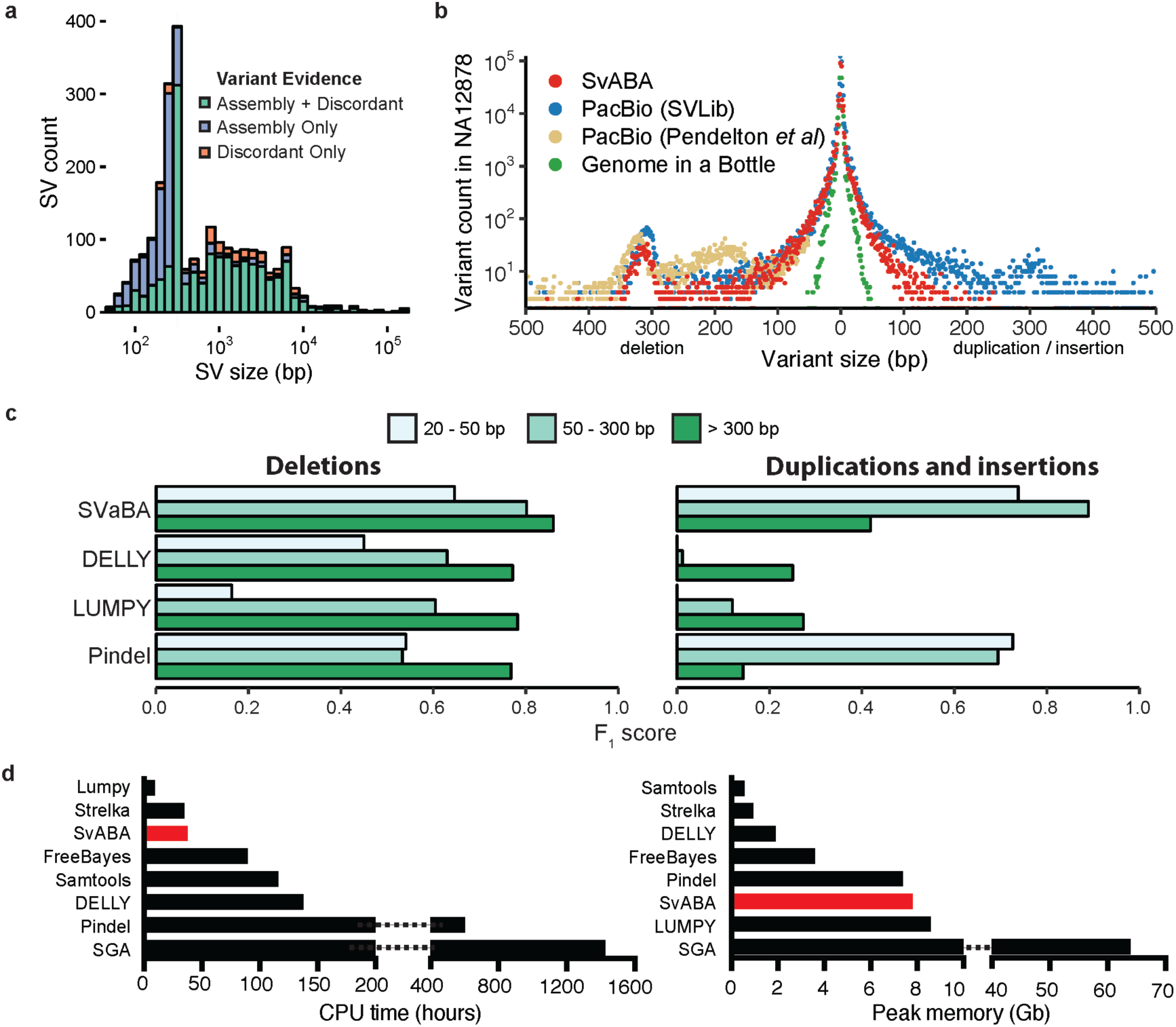
Detection of SVs and indels in the NA12878 human genome. **a**) The number of SV events and the types of supporting evidence used by SvABA for detecting SV events of different lengths (indel variants not shown). SVs are detected through realignment of assembly contigs (purple), discordant read clusters (orange), or a combination of both (green). SVs with shorter lengths than the average size of the sequencing fragments are identified almost exclusively through assembly and realignment. **b**) The length distributions of indels and small SVs in NA12878 determined from different sequencing and analytical technologies: 151 base paired-end Illumina sequencing by SvABA (red), PacBio-assemblies from Pendleton *et al*(Pendleton et al. 2015) and SVlib, PacBio-assembly and raw PacBio-read SV calls from Pendleton *et al*(Pendleton et al. 2015) (only deletions > 50 bp shown; light brown), and the indel callset of the Genome in a Bottle consortium(Zook et al. 2014) (green). **c**) Comparison of detection accuracy of SvABA, LUMPY, DELLY and Pindel for deletions (left), and for insertions/duplications (right), across three different length regimes in NA12878. The F_1_ score is a combined measure of precision and recall, and was calculated using the PacBio-assemblies and GIAB as a truth set. **d)** Total CPU and peak memory usage for several indel and SV detection tools applied to a single 33x human genome. SGA CPU and memory usage was estimated using published data (Simpson and Durbin 2012).

We next evaluated the performance of SvABA using two truth sets: indels from the Genome In a Bottle (GIAB) integrated callset (Zook et al. 2014), and SVs and indels from SVlib operating on contigs from a recent whole genome *de novo* assembly of Pacific Biociences (PacBio) reads by Pendleton *et al* (Pendleton et al. 2015). The GIAB integrated callset provides high-confidence short indels by combining 13 different short-read variant callers applied to short reads sequencing data, but is not designed to detect larger variants. The substantial length of the PacBio-assembly contigs makes them an appropriate platform for detecting larger SVs as well as short indels. We reasoned that GIAB and PacBio sequencing would provide complementary evidence and cover the full range of variants sizes called by SvABA.

To generate calls from the PacBio-assemblies, we aligned the contigs to the reference with BWA-MEM and used SVlib to parse the gapped and multi-part alignments to produce the SV and indel variants. We then genotyped the variants by realigning the 151 base sequence reads to the assembly contigs. After masking regions with low sequence mappability, SVlib on the PacBio-assemblies identified 4,482 deletions, 4,278 duplications, 488 inversions, 235,608 small deletions (≤ 50 bp) and 205,791 small insertions (≤ 50 bp) (**Supplementary Data 2**). Of the indels, 131,989 (29.9%) overlapped a variant in the GIAB callset. PacBio-assembly indels that did not overlap a GIAB variant were contained in regions of lower mappability (p < 1e-16, Wilcoxon-test) and had a higher mean size (GIAB: 3.03 bp, PacBio: 3.99 bp, p < 1e-16 Wilcoxon-test). This is consistent with other reports showing long read PacBio sequences are more powered to capture variation in repetitive regions and across a wide range of sizes (Huddleston et al. 2016). We additionally included in the PacBio truth set the 7,959 deletions that were identified by Pendleton *et al* (Pendleton et al. 2015) using the both raw PacBio reads and assemblies and the PBHoney detection tool (English et al. 2014).

SvABA detected a significantly greater number of small SVs and indels than are present in the GIAB set, and exhibited a variant size distribution very similar to that of the PacBio-assembly variants (Fig. 2b). Among the short indels (≤ 50 bp) called by SvABA, 72.2% were contained in either the GIAB or PacBio-assembly callset. The proportion of SvABA indels in the GIAB set was strongly size dependent, with indels < 10 bp being nearly 4 times more likely to be present in the GIAB set than indels > 40 bp (**Supplementary Fig. 5**). Private SvABA calls were significantly enriched for heterozygous variants (p < 0.001, Fisher’s exact), which in many cases appeared to be at sites with only one allele represented by the PacBio-assembly.

We next used the PacBio-assembly calls as a truth set to compare the detection performance of SvABA versus three SV callers: LUMPY, Pindel and DELLY. We selected these callers because of their wide use in the field, and because of their different approaches to SV detection, including combinations of inference from discordant reads, split read alignments, and local sequence realignment within each tool. For each of three different size regimes (20-50 bp, 51-300 bp and 300+ bp), we calculated the F_1_ score, which is a combined measure of the sensitivity and specificity.

SvABA was sensitive to events across the full range of sizes (Fig. 2c), and exhibited the greatest sensitivity for duplication and insertion variants (Table 1) For medium-sized duplication events between 50 and 300 bp, SvABA detected 1.6-fold as many true events as the next most sensitive method, Pindel. The sensitivity gain for medium-sized duplications was more pronounced when compared with DELLY (163-fold) and LUMPY (15-fold). For small deletions (20 - 50 bp), Pindel and SvABA exhibited the greatest overall sensitivity of the four, with Pindel identifying 1.15-fold more variants than SvABA, but with a 2.1-fold higher unvalidated rate. For large deletions, DELLY and SvABA exhibited the highest sensitivity, with DELLY identifying 1.06- fold more deletions, but with a 3.7-fold higher unvalidated rate.

**Table 1:**
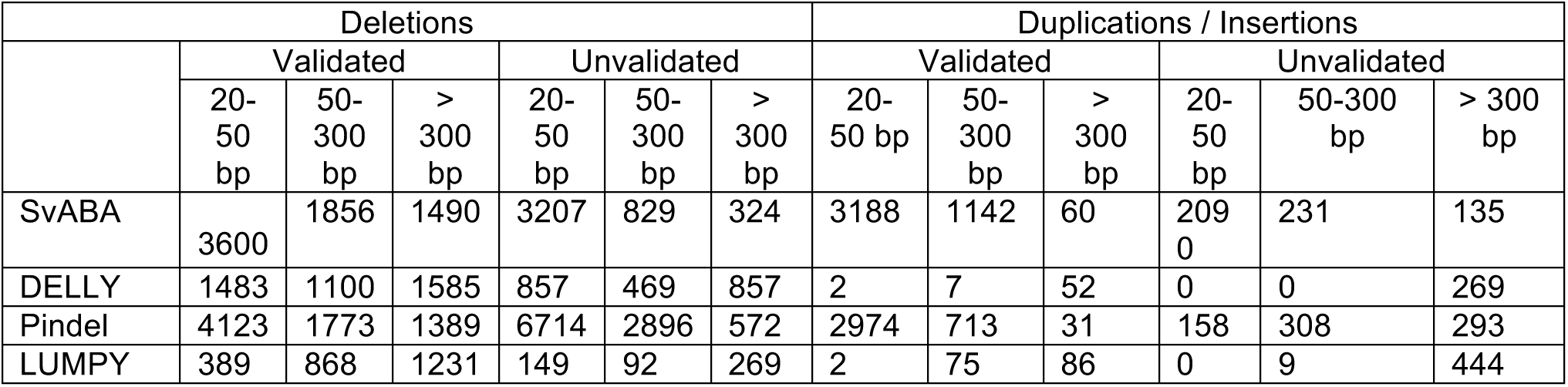
SV and large indel detection in NA12878, validated against PacBio assemblies and Genome in a Bottle indels

SvABA was able to achieve these results without requiring extensive CPU or RAM allocations. A major obstacle to assembly-based variant detection has been the computational requirements needed for *de novo* assembly. However, SvABA assembled the NA12878 data in 3,150 CPU minutes using 7.7 Gb of memory, which was primarily used to store the indexed reference genome. SvABA required fewer CPU resources than all other detection tools except LUMPY (Fig. 2d). With the native parallelization in SvABA, we distributed the compute over 12 cores to call variants in just over 5 wall-time hours (313 minutes). We further tested the ability of SvABA to perform in an ultra-highly parallel computing environment and were able to call variants in 144 minutes on a single 33x BAM using 30 CPU cores.

### Validation of SvABA using an in silico tumor model

Detection of somatic SVs and indels in cancer poses significant challenges beyond those faced in detecting germline events. Somatic rearrangements involve a higher rate of large and inter-chromosomal events (Yang et al. 2013), and are often clustered tightly with other rearrangements as part of complex events like chromothripsis (Stephens et al. 2011). Rearrangements in cancer are frequently repaired by non-homologous end joining (NHEJ) (Yang et al. 2013), a process which can introduce novel base pairs at rearrangement junctions during the repair process. Somatic variants must also be distinguished from the background of germline variation; the challenge is further amplified by wide variability in relative amounts of normal and tumor cells in tissue samples. We therefore set out to separately validate SvABA as a tool for detecting somatic SVs and indels in a way that accounts for these challenges.

Due to the difficulty of obtaining gold-standard truth sets for somatic SVs from real data, we opted to use SVlib to generate an *in silico* tumor-normal pair that more closely reflected the unique challenges of SV detection in the cancer genome. The frequency of somatic copy-number events is known to be inversely proportional to the length of the event (Zack et al. 2013), and we recapitulated this in our simulation by creating a range of indels and rearrangements that mirrored this length distribution (**Supplementary Fig. 6**). We also spiked in 2000 short (<= 10bp) indels to better reflect the high indel rates seen in real tumors. Our *in silico* tumor contained 49,995 heterozygous SVs and indels, including 2,741 duplications, 5,444 deletions, 21,334 large (> 2,000 bp) intra-chromosomal rearrangements, 9,037 small deletions (< 50 bp) and 8,918 small insertions. To model the novel junction insertions of NHEJ, we supplied half of the junctions with novel 1-20 bp insertions. We then simulated 101 base paired-end reads from the virtual tumors at coverage of 10x and a mean insert size of 250 bp using ART (Huang et al. 2012). To simulate a sample with low tumor purity, we mixed the simulated tumor reads with real reads from the HCC1143BL lymphoblastic normal cell line at 30x coverage. Both the simulated read pairs and HCC1143BL reads were aligned to the reference with BWA-MEM (see Methods for data availability). We then called somatic variants on our *in silico* tumor using SvABA, FreeBayes, Strelka, DELLY, LUMPY and Pindel.

SvABA reached the greatest overall sensitivity among the six methods, detecting 87.4% of all variants and achieved the highest F_1_ score for each size range of variants (Table 2). SvABA was modestly more sensitive overall than FreeBayes and Strelka for indels (1.1-fold increase), but captured a substantially higher number of indels greater than 20 bp (Fig. 3a). Pindel similarly achieved relatively broad coverage across different sizes, but was less sensitive to insertion variants than SvABA, and required substantial filtering across all variant sizes in order to achieve similar false positive rates as the other methods. DELLY and LUMPY performed similarly for both medium-sized and large SVs, with LUMPY achieving the lower false positive rate. However, SvABA substantially improved detection (1.6-fold increase) relative to DELLY and LUMPY for medium-sized SVs at similar false positive rates. For large variants (> 300 bp), SvABA, DELLY and LUMPY exhibited largely similar performance, with SvABA achieving the highest sensitivity by a small margin.

**Table 2:**
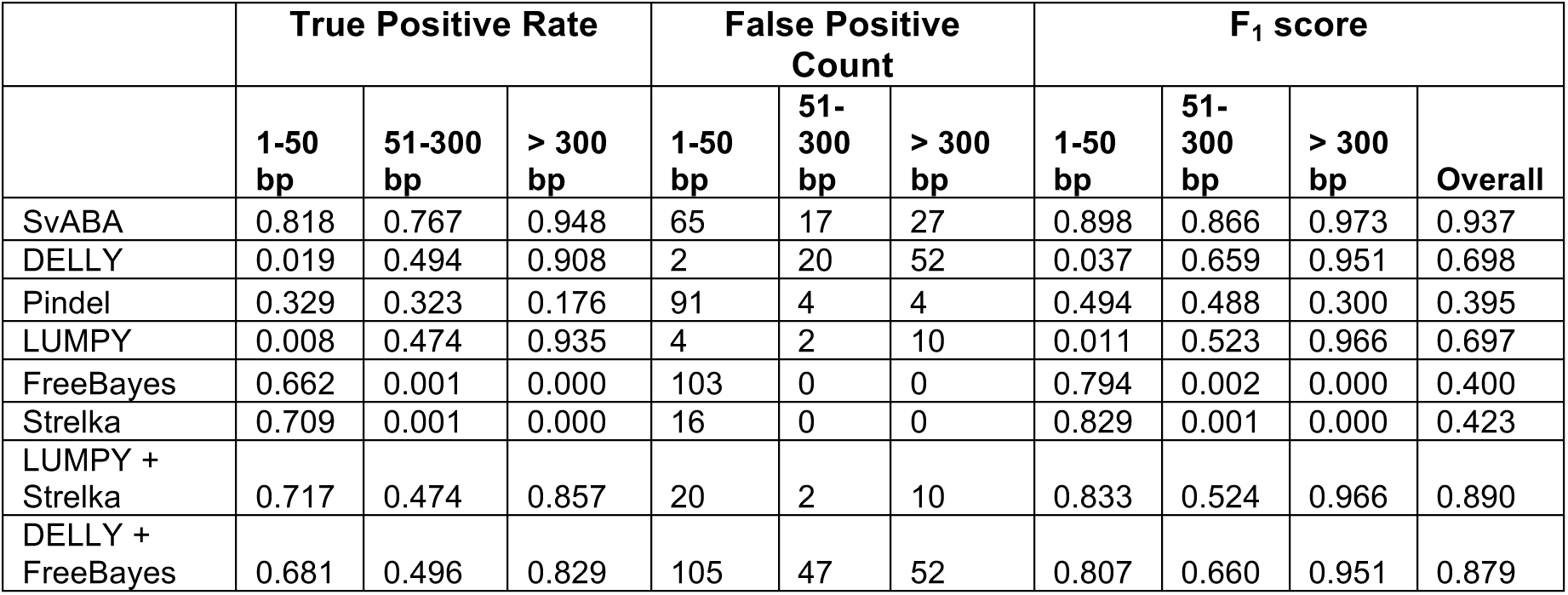
Somatic indel and SV detection performance using an *in silico* tumor genome

|  | True Positive Rate |  |  | False Positive Count |  |  | F<sub>1</sub> score |  |  |  |
| --- | --- | --- | --- | --- | --- | --- | --- | --- | --- | --- |
|  | 1-50 bp | 51-300 bp | > 300 bp | 1-50 bp | 51-300 bp | > 300 bp | 1-50 bp | 51-300 bp | > 300 bp | Overall |
| SvABA | 0.818 | 0.767 | 0.948 | 65 | 17 | 27 | 0.898 | 0.866 | 0.973 | 0.937 |
| DELLY | 0.019 | 0.494 | 0.908 | 2 | 20 | 52 | 0.037 | 0.659 | 0.951 | 0.698 |
| Pindel | 0.329 | 0.323 | 0.176 | 91 | 4 | 4 | 0.494 | 0.488 | 0.300 | 0.395 |
| LUMPY | 0.008 | 0.474 | 0.935 | 4 | 2 | 10 | 0.011 | 0.523 | 0.966 | 0.697 |
| FreeBayes | 0.662 | 0.001 | 0.000 | 103 | 0 | 0 | 0.794 | 0.002 | 0.000 | 0.400 |
| Strelka | 0.709 | 0.001 | 0.000 | 16 | 0 | 0 | 0.829 | 0.001 | 0.000 | 0.423 |
| LUMPY + Strelka | 0.717 | 0.474 | 0.857 | 20 | 2 | 10 | 0.833 | 0.524 | 0.966 | 0.890 |
| DELLY + FreeBayes | 0.681 | 0.496 | 0.829 | 105 | 47 | 52 | 0.807 | 0.660 | 0.951 | 0.879 |

**Figure 3:**
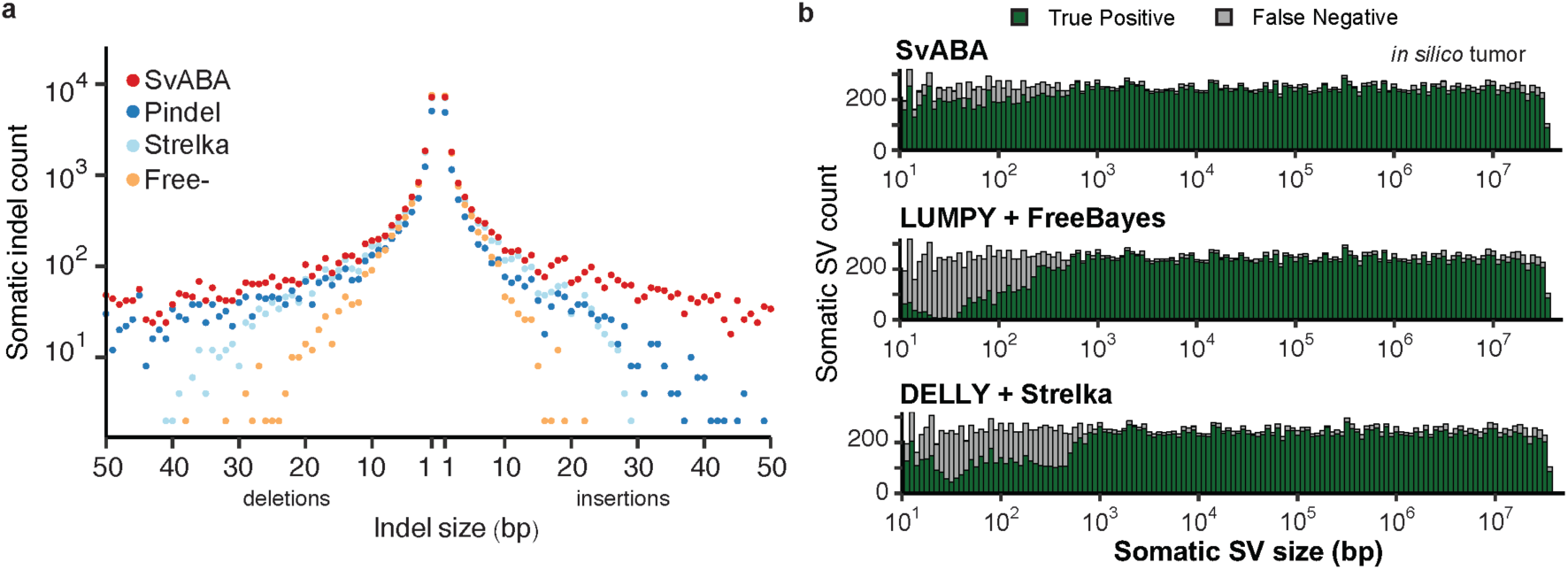
Benchmarking somatic variants with an *in silico* tumor. **a)** True positive counts for indel calling (y-axis) as a function of variant size (x-axis) for SvABA (red), Pindel (green), FreeBayes (orange), and Strelka (purple). All callers achieved similar sensitivities for small somatic indels, while SvABA maintained high-sensitivity for larger indels (> 10 bp) for both insertions and deletions. **b)** Stacked bar-chart of the number of SVs detected across all SV types (y-axis) as a function of variant size (x-axis). SvABA (top) maintained sensitivity across variants of all sizes. Combining calls from a dedicated indel and SV caller (middle: LUMPY and FreeBayes, or bottom: DELLY and Strelka) improved overall sensitivity, but still left a gap for medium-sized SVs.

We further considered how the detection performance of SvABA would compare with the combined performance of using both an indel caller and an SV caller together on the same data. We paired the calls from DELLY and FreeBayes together, and the calls from LUMPY and Strelka together, to create two examples of using combined callsets to evaluate variants. The two combined callsets reached similar performance as measured by the F_1_ score (LUMPY + FreeBayes: 0.890, DELLY + Strelka: 0.879), but both were lower than SvABA (0.937). In both cases, the combined callers differed most greatly from SvABA for variants between 50 bp and 300 bp, where SvABA detected 1.5-fold more variants than either combined approach (Fig. 3b).

### Detection performance in comparison with whole-genome de novo assemblies

We next evaluated the performance of SvABA in real data from a human tumor. We used data from two separate library preparation and sequencing strategies in the HCC1143 breast cancer cell line and its paired lymphoblastic normal HCC1143BL. The first dataset was sequenced from libraries prepared with a standard Illumina PCR amplification step and 101 base paired-end reads. The second dataset was sequenced from libraries prepared without the PCR amplification and using 250 base paired-end Illumina reads (Kozarewa et al. 2009).

To provide an alternative computational approach using the 250 base PCR-free reads, we performed whole-genome *de novo* sequence assembly using Discovar *de novo*, the whole-genome *de novo* assembly successor to DISCOVAR (Weisenfeld et al. 2014). Discovar *de novo* is specifically designed to assemble 250 base read data, and contig generation is unbiased by alignment issues. We extracted SVs and indels from the Discovar *de novo* assemblies by aligning the Discovar *de novo* contigs to the reference with BWA-MEM, and then parsed the gapped and multi-part alignments with SVlib. We further distinguished somatic and germline variants by realigning the 250 base reads from both HCC1143 and HCC1143BL within SVlib to the Discovar *de novo* contigs, and evaluated the tumor and normal support for each variant.

The three callsets exhibited substantial overlap, with the main difference being an increased sensitivity from the longer read lengths and a smaller sensitivity gain from the whole-genome *de novo* assembly as compared to genome-wide local assemblies (**Supplementary Table 1;** Fig. 4a). Discovar detected the highest number of somatic variants (1,538), followed by SvABA on the 250 base reads (1,409) and then SvABA on the 101 base reads (1,016). With the standard 101 base reads, SvABA achieved a high specificity, with 92.8% of variants being rediscovered in the 250 base reads. When comparing SvABA and Discovar calls from the same dataset (250 base PCR-free reads), SvABA detected 69.9% of Discovar variants; conversely, 76.3% of SvABA variants were present in the Discovar results. Relaxing the read-support threshold for the Discovar calls increased the support for 250 base SvABA calls to 89.1%.

**Figure 4:**
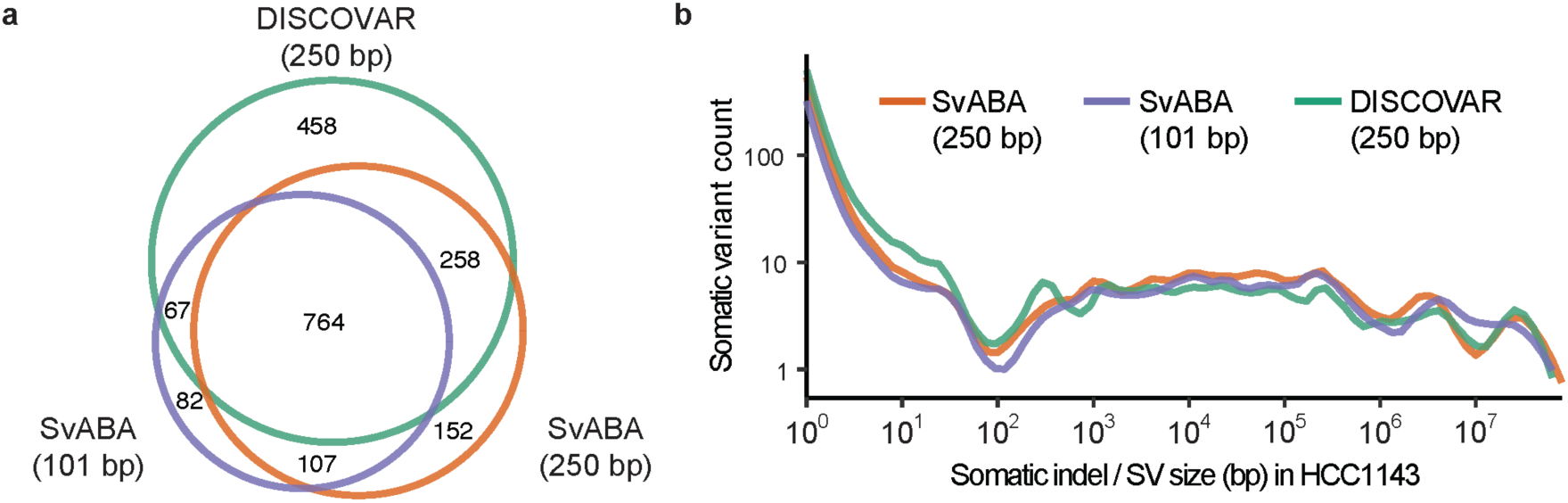
Somatic variant detection in the HCC1143 breast cancer cell line using different sequencing and informatics approaches. **a)** Comparison of combined somatic SV and indel detection in HCC1143 using: local assembly using SvABA with 101 base read paired-end reads (purple), SvABA with 250 base paired-end PCR-free reads (orange), or global assembly using Discovar *de novo* on 250 base paired-end PCR free reads and SVlib to extract variants (green). Somatic variant counts (y-axis) for Discovar *de novo* (250 base read PCR-free reads;; green) and SvABA using 101 base read (purple) or 250 base read PCR-free reads (orange), as a function of variant size (x-axis). All methods have similar sensitivities across different sizes, except Discovar *de novo* was more sensitive to short indels in simple repeats.

Variants detected with Discovar and SvABA show nearly identical size distributions (Fig4b). The events that were discovered by Discovar but not SvABA with either dataset were highly enriched for events occurring near centromeres (p < 0.01, Fisher’s exact) and in simple repeats (p < 0.01). This is consistent with an improved ability of long-reads and global *de novo* assembly to identify variants in regions of the genome that are difficult to align to, likely resulting from the improved alignment quality and reduced alignment ambiguity afforded by long sequences.

#### Somatic rearrangements frequently involve short templated-sequence insertion junctions

Complex events are increasingly recognized to be prevalent in both germline and cancer genomes (Chiang et al. 2012;; Stephens et al. 2011). However, their detection is complicated when neighboring breakpoints are separated by distances on the order of the read length or greater, due to the difficulty in aligning short reads covering such divergent sequences. We hypothesized that assembly-based methods might have superior sensitivity for such events Assembly contig generation can occur directly from clipped and unmapped alignments, and builds consensus sequences regardless of breakpoint complexity. Indeed, while inspecting the Discovar and SvABA contigs from HCC1143, we identified multiple contigs that contained three or more sequences with high-quality alignments to disparate genomic loci, supporting putative complex events with multiple neighboring breakpoints (Fig 5a).

**Figure 5:**
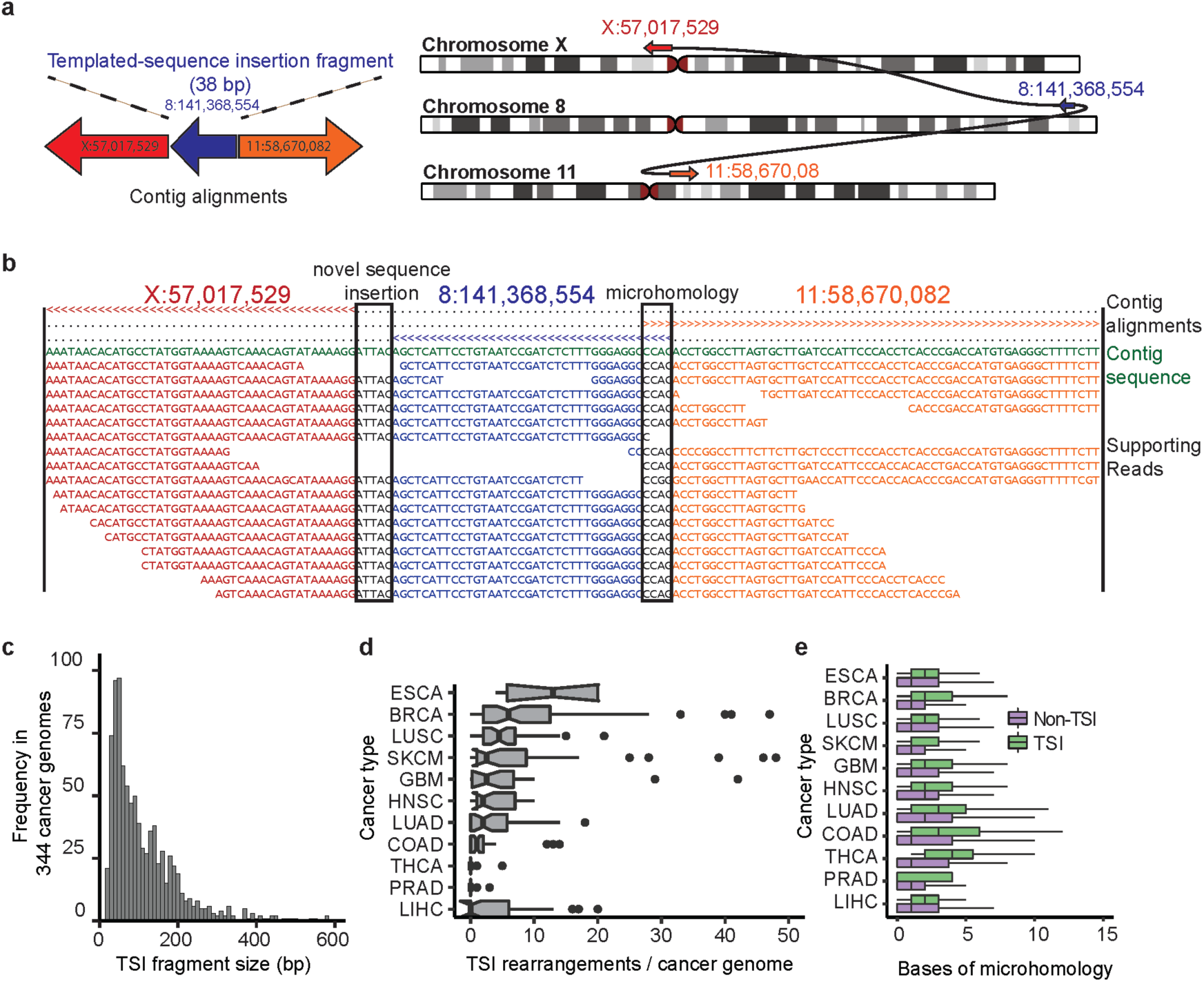
SvABA identifies rearrangements with templated-sequence insertions (TSI) derived from distant genomic loci. **a)** Somatic rearrangement between chrX and chr11 in HCC1143 containing a 38 bp fragment of chr8. TSI rearrangements are identified by assembly contigs that have multiple non-overlapping alignments to the reference. The direction of the arrows represents the strand that the contig fragment was aligned to (right-facing is forward strand). **b)** Partial view of the contig from (a) showing the multiple alignments of the contig to the reference and the read-to-contig alignments. The top three lines indicate which bp of the contig each of the three BWA-MEM alignments covers (> is forward strand alignment, < is reverse strand alignment). The first two alignments indicate an insertion of 5 bp of novel sequence at the first junction (left), and the second two indicate 4 bp of microhomology at the second junction (right). The middle alignment supports the TSI fragment. These plots are automatically generated by SvABA for each variant (in the *.alignments.txt.gz file). **c)** TSI fragment lengths from somatic rearrangements across 344 cancer genomes (mean 86 bp). **d)** Prevalence of TSI rearrangements (x-axis) across 11 tumor types (y-axis). ESAD: esophageal cancer, BRCA: breast cancer, LUSC: lung squamous cell carcinoma, SKCM: melanoma, GBM: glioblastoma multiforme, HNSC: head and neck squamous cell carcinoma, LUAD: lung adenocarcinoma, COAD: colorectal adenocarcinoma, THCA: thyroid carcinoma, PRAD: prostate adenocarcinoma, LIHC: hepatocellular carcinoma. **e)** Bases of breakpoint microhomology (x-axis) for different cancer types (y-axis) for somatic TSI rearrangements (green) and somatic non-TSI rearrangements (red). The TSI rearrangements have a significantly higher degree of breakpoint microhomology than their non-TSI counterparts across all tumor types.

We therefore investigated whether SvABA could systemically discover complex events from the HCC1143 101 base read data. Among the SvABA contigs generated from the 101 base read data, we found eight contigs with high-quality multi-part alignments (**Supplementary Table 2**). These complex contigs were well supported throughout their length by sequence reads, and we found no significant difference in the number of breakpoint-supporting reads between simple and complex events (mean 52.4 in multi, 50.4 simple, p = 0.69, t-test). There was no significant difference in the mapping quality of the alignments between simple and complex rearrangements (mean 55.1 in multi, 58.5 in simple, p = 0.17, t-test). We concluded that these sequences represented true rearrangements containing templated-sequence insertions (TSI) copied from distant loci. Rearrangements involving templated-insertions have been described in the germline at the junctions of larger complex rearrangements (Liu et al. 2011) and in cancer in the context of chromothripsis events (Zhang et al. 2015), but have not been otherwise extensively described in cancer genomes. Therefore, we wished to specifically validate these events in HCC1143 and evaluate their prevalence across other cancers.

Among the eight contigs directly supporting a TSI rearrangement, seven were directly supported by a Discovar contig from the 250 base read data. To confirm that the multi-part alignments did not have a more parsimonious non-TSI alignment, we realigned the seven Discovar-supported contigs using BLAT (Kent 2002). BLAT confirmed the alignments from BWA-MEM for each of the alignment fragments outside of the TSI fragments. BLAT further supported a non-local alignment for each of the TSI fragments, with some ambiguity as to the originating location for the shortest insertion fragments (**Supplementary Table 2**).

The TSIs from single contigs were short (median 56.5 bp), and we hypothesized that additional TSI rearrangements with longer insertions could be found by clustering together chains of rearrangements from multiple contigs, with breakpoints separated by fewer than 1,000 bp. This yielded 30 separate rearrangement clusters, including those from the single-contig rearrangements (**Supplementary Table 3; Supplementary Figure 7**). The median fragment size across these clusters was 185 bp. Nine of the clusters involved contiguous chains of multiple TSIs, including cluster 25 that contained 5 contiguous TSI fragments. This cluster was supported throughout by a Discovar contig.

We also confirmed the predicted sequences for eight TSI junctions by performing PCR spanning the junctions, including ones identified by single and multiple contigs (**Supplementary Fig. 8, Supplementary Table 4**). The rearrangements were validated as somatic, as none of these junctions were detected by PCR in the HCC1143BL normal cell line. These results confirm both the presence of these rearrangements, and provide direct evidence that the multiple breakpoints are present on the same allele. Based on the close proximity of the breakpoints and the results of our validation, it is likely that the remaining clusters represent rearrangements from the same allele. Long-read sequencing would be required to systematically validate genome-wide the phasing of clustered rearrangements and rearrangements with TSI fragments longer than the library fragment size.

These short TSIs (on the order of a read length) are not restricted to the HCC1143 cell line, but rather appear across a range of cancers. To test whether TSI junctions could be discovered across a range of tumor types, we ran SvABA on 344 TCGA whole-genome tumor-normal pairs (**Supplementary Table 5**). SvABA called 47,965 rearrangements, including 2,124 rearrangements (4.4% of all somatic rearrangements) with evidence of TSI junctions. The mean TSI size was 89.0 and ranged between 21 bp and 1082 bp (Fig. 5b). The number of TSIs per tumor was highly correlated with the number of simple events (R^2^ = 0.38). Esophageal carcinomas exhibited the highest rate of TSI rearrangements, with a median of 13 per sample, followed by breast with six and lung squamous carcinoma with 4.5 (Fig. 5d). These tumors also exhibited the highest median numbers of rearrangements with non-TSI junctions (esophageal 192, breast 226.5, lung squamous 99.5 per sample). Finally, although somatic rearrangements were largely mediated by non-homologous repair, somatic TSI events contained over twice the amount of microhomology than non-TSI somatic junctions (2.82 bp in TSI, 1.28 in non-TSI, p < 0.001, t-test; Fig. 5e). Although somatic rearrangements are thought to be largely mediated by non-homologous repair, the subtle but significant difference in the length of homology between TSI and non-TSI junctions may reflect different underlying mechanisms generating these events.

The prevalence of TSIs in the cancer genome suggests that they may underlie oncogenic events. As an example of an oncogenic driver alteration formed in part by TSI events, we identified a focal amplification in a glioblastoma of the EGFR receptor tyrosine-kinase containing 52 non-TSI junctions and seven TSI junctions (**Supplementary Fig. 9**). For many of the TSI junctions, the supporting reads were comprised of primarily reads that were initially unmapped, and thus rescued by the local assemblies.

#### SvABA identifies sites of viral integration

Insertion of DNA sequences from viral and bacterial pathogens represents an important mechanism of oncogenesis, but these events will often go undetected because pathogen sequences typically do not align to the reference genome. We hypothesized that an assembly-based approach could provide greater sensitivity to detection of these events because of our systematic analysis of unmapped reads with mapped pair-mates. To detect viral insertions, we built SvABA to accept an alternate reference genome of non-human sequences, in addition to the primary human reference. In this mode, locally assembled contigs that significantly align to the alternative reference are then emitted in the VCF as fusions between the human and non-human genome coordinates.

As a proof of principle of how this might be used, we ran SvABA on 16 head and neck carcinomas using the RefSeq viral sequence database as an alternative genome and looked for evidence of integration of the human papillomavirus (HPV). HPV is a known oncovirus in head and neck cancer, and is known to integrate into the genome of tumor cells (Parfenov et al. 2014). SvABA identified 16 breakpoints across 7 samples where viral sequences were fused with genomic DNA (**Supplementary Table 6**). All of the viral junctions involved integration of HPV type 16 into the genome, and each of the insertion sites validated by comparison with Parfenov *et al* (Parfenov et al. 2014).

#### Medium-sized SVs as driver alterations in cancer

We next examined the contribution of small and medium-sized (< 500 bp) insertion, deletion and duplication variants as potential driver events in cancer. Using the Cancer Gene Consensus list of cancer genes, we evaluated the relative burden of somatic indels and SVs in the exons of known cancer genes versus non-cancer genes. Across all size regimes, small and medium-sized variants were significantly enriched in cancer genes (p < 0.01, Fisher’s exact, Fig. 6a). We identified 19 indels and SVs with sizes between 21 bp and 445 bp that altered the exon of a known cancer gene (**Supplementary Table 7**). Most of these (13 of 19) were smaller than 50 bp.

**Figure 6:**
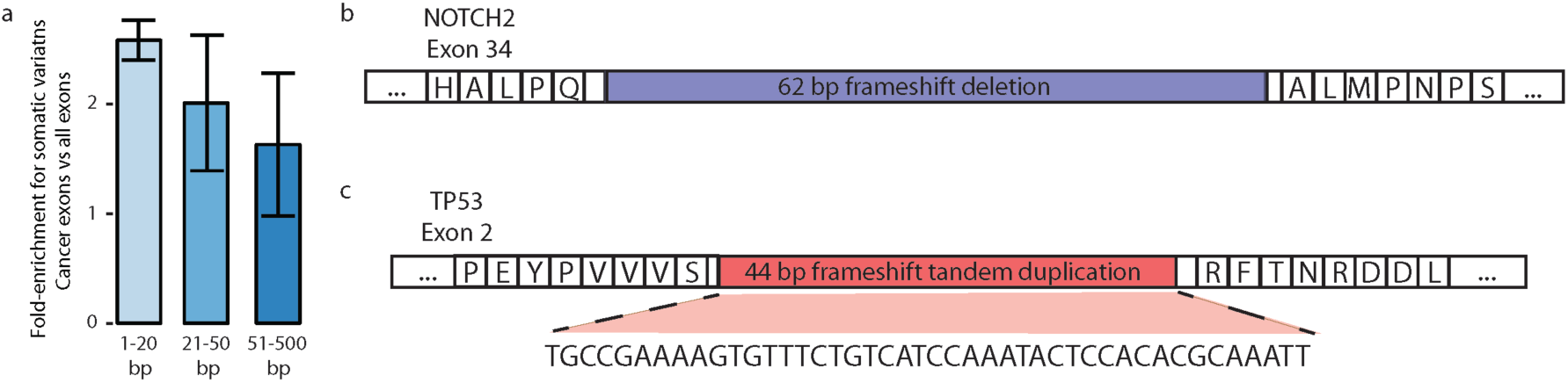
Contribution of medium-sized SVs to disruption of known driver genes in cancer. **a)** Fold-enrichment (y-axis) for breakpoints from SVs and indels of three different size regimes (1-20 bp, 20-50 bp, 51-500 bp) occurring inside exons of known cancer genes versus from all exons. **b**) A frameshift deletion of 62 bp in exon 34 of NOTCH2 in a breast adenocarcinoma (TCGA-AO-A0J6-01). The partial amino acid sequence of exon 34 is shown for reference. **c)** A likely loss-of-function 44 bp frameshift tandem duplication in exon 2 of *TP53* in a lung squamous cell carcinoma (TCGA-68-7755).

Calling these 21-500 bp SVs may be necessary for accurate genotyping of cancer genes. For example, we identified a 62 bp frameshift deletion in exon 34 of *NOTCH2* in a breast adenocarcinoma (Fig. 6b). C-terminal *NOTCH2* alterations have been found to be recurrent in B cell lymphomas, and lead to a gain-of-function product (Lee et al. 2009). Increased *NOTCH2* expression is also associated with decreased survival in breast cancer patients (Parr et al. 2004). We also identified a 44 bp tandem duplication in exon 2 of the *TP53* tumor suppressor gene in a lung squamous cell carcinoma (Fig. 6c), indicating loss of *TP53* function. Based on these findings, we expect that additional driver alterations in this size regime could be discovered in future cancer genome analyses by incorporating genome-wide assembly-based detection methods like SvABA.

### Discussion

We found that genome-wide local assembly exhibits broad sensitivity for indel and SV detection across arange of variant sizes. Our assembly-based approach was particularly sensitive for variants between 20 and 300 bp, and robustly identified complex rearrangement junctions containing templated-sequence insertions (TSI) and sites of viral integration. The ability to detect such a broad range of variants within a single framework represents an important advance towards achieving complete characterization of genomes from short-read sequencing data. As a demonstration of how our approach may be used to identify novel biologically relevant variants, we discovered several cases where complex rearrangement junctions and small SVs contributed towards driver events in cancer.

Despite being a primarily assembly-based detection tool, we found that integrating both assembly and alignment signals improved the overall detection performance relative to either alone. Integrative approaches that combine multiple read signals in one inference framework, notably LUMPY and DELLY, have been previously shown to boost detection performance over any single approach. With SvABA, the addition of genome-wide local assembly provided an important signal for discovering medium-sized and complex variants, while providing additional support for large variants with more robust discordant read evidence. In addition to implementing assembly-driven variant detection, SvABA provides a number of improvements over alignment-based approaches, including realignment of discordant reads and pair-mate region lookups to boost the read support without requiring any BAM preprocessing.

Our implementation of a genome-wide assembly approach for SV detection builds on the work described by TIGRA and others, which used targeted assemblies to improve specificity for SVs within the 1000 Genomes Project (Sudmant et al. 2015). The primary contribution of SvABA is to apply local assembly genome-wide instead of to a smaller set of regions (e.g. those nominated through discordant mate clustering or soft-clipped alignment). To overcome the computational challenges of this approach, we first developed the SeqLib library to provide SvABA with extremely efficient access to BAM files and in-memory BWA indexing and alignment capabilities (Wala and Beroukhim 2016). We also refactored SGA to provide for rapid in-memory microassemblies. Our resulting method unites alignment, assembly and variant calling within a single C++ program, making possible the genome-wide local assembly approach with a single command and pass through the BAM file. The local-assembly problem is also inheritably parallelizable, and we built SvABA with native parallelization to any number of CPU cores. This allows for genome-wide discovery within only a few hours, and is as easy to use as the multithreaded capabilities of tools like BWA.

There are limitations to a local assembly-based approach. SvABA relies strongly on having sufficient variant reads to build an assembly contig. For low-coverage genomes or for highly impure tumor samples, the number of reads may not be sufficient to provide for robust assemblies. LUMPY, DELLY and Meerkat, among other tools, have been specifically tuned to detect variants in genomes with low coverage, and may be more sensitive than SvABA for low-coverage data. SvABA also relies on the approximately correct alignment of at least one read in a pair. Read pairs where both ends are unaligned or incorrectly mapped will not be accurately evaluated by SvABA, but may be correctly incorporated by genome-wide *de novo* assembly. As such, while SvABA does not rely on any one particular alignment method, the initial alignments should be robust enough to place a sufficient number of read pairs approximately near the breakpoints. Indeed, we observed an increase in sensitivity with whole-genome *de novo* assemblies using Discovar *de novo*, although its computational requirements make it currently infeasible to apply to large cohorts. Finally, SvABA uses BWA-MEM for contig alignment and variant calling. Though BWA-MEM enables very rapid alignment of long sequences to the reference, it may be more appropriate to use a more sensitive algorithm for highly divergent queries.

Even with improved informatics approaches like *de novo* assembly, SV detection in short-read sequencing data is ultimately limited by the read lengths – improved detection requires technologies that produce long-range information that can fully capture variation in repetitive regions or from highly complex rearrangements. We developed SVlib to call indels and SVs from PacBio-assemblies, and other approaches such as PBHoney (English et al. 2014), MultiBreak-SV (Ritz et al. 2014) and hySA (Fan et al. 2017). have also been developed for extracting larger SVs from long-sequences. Alternatively, short-read sequencing may be combined with barcode libraries that produce linked-reads (e.g. from 10X Genomics). Linked reads are also particularly useful for long-range haplotyping of detected variants (Zheng et al. 2016).

We found that each of the tools we benchmarked against provided excellent calls within their targeted size regimes. For instance, we found Pindel to be quite sensitive for small deletion variants, while LUMPY and DELLY achieved highly accurate detection of larger SVs. With SvABA, we have provided a single method that achieves high accuracy in both of these size regimes, and additionally covers the gap in between short indels and larger SVs. An alternative approach to using a single caller is to integrate results from multiples methods covering a variety of detection approaches, as was done by the 1000 Genomes Project to achieve broad SV sensitivity in their recent survey of 2,504 human genomes (Sudmant et al. 2015). Integrating callsets is also valuable for increasing specificity, and we expect that SvABA will be a useful addition to large-scale sequencing efforts and consortium SV calling. For instance, SvABA has been recently used to generate somatic variant calls from 2,961 cancer whole-genomes as part of the International Cancer Genome Consortium (ICGC). We expect that SvABA’s low computational burden and ease of running, and hence low cost to operate, coupled with its broad sensitivity and applicability to germline or cancer genomes, will continue to make it a practical and suitable tool for such large-scale analyses.

## Methods

### Read retrieval from local assembly windows

SvABA extracts the following reads by default: alignments with high-quality clipped bases, discordant reads, unmapped reads, reads with unmapped pair-mates, and reads with deletions or insertions in the CIGAR string. Reads that are marked as PCR duplicates, failed QC reads, and reads with homopolymer repeats > 20 bp are removed before assembly. SvABA additionally considers a read a duplicate if it has the same sequence, alignment position and pair-mate position as another read.

To avoid computational bottlenecks associated with ultra-high coverage regions, extracted reads are subsampled to keep coverage below a user-specified level (default=100). SvABA also allows users to exclude pre-specified regions. A recommended exclusion interval file that masks centromeres and alpha satellite regions is provided for hg19 at https://data.broadinstitute.org/snowman/svaba_exclusions.bed. Targeted assemblies can be performed by providing a BED file or coordinate string (e.g. 1:1,000,000-2,000,000).

### Read trimming and error correction

To reduce assembly errors from low-quality bases, reads are trimmed at their 3’ and 5’ ends prior to assembly. Starting from each end of the read, bases are removed until a high quality (scaled-base-quality score > 4) is identified. Reads trimmed to fewer than 30 bases are removed. Reads with a clipped alignment size and insert-size consistent with sequencing into the sequencing adapter are excluded.

SvABA wraps two error-correction strategies that can be toggled by the user at run-time: Bloom-filter correction BFC (Li 2015), and a *k*-mer tracking method as implemented in SGA(Simpson and Durbin 2012). BFC is the default error correction method because of its improved speed over the k-mer strategy implemented in SGA, and its ability to error-correct both gapped alignments and single nucleotide errors. For the *k*-mer method, SvABA uses a *k*-mer size of 31, and accepts *k*-mers with three or more instances as representing a true sequence. SvABA can optionally emit a FASTA file of error-corrected sequences. For both types of error correction, all of the reads from the assembly window, including non-variant reads, are used to train the error corrector.

### Discordant read realignment and clustering

The insert-size distribution for each read group is estimated from a sample of five million reads. Only read-pairs with a forward-reverse pair orientation are used for estimating the insert size. To exclude read-pairs with unusually large or small insert sizes that likely represent misalignment or true variants, the largest and smallest 5% of insert-sizes are removed from the insert-size estimation. Reads with non-standard pair-orientations or outlier insert-sizes greater than four standard deviations from the expected insert-size for that read group are considered candidate discordant reads.

Due to the difficulty of aligning reads in non-unique regions of the genome, most discordant reads can be attributed to mapping artifacts rather than true variation. To reduce the effect of alignment artifacts on generating false-positive variant calls, candidate discordant reads are realigned on-the-fly with BWA-MEM to the reference genome. Candidate discordant reads with an available non-discordant alignment of greater than 70% of the maximum alignment score are removed from discordant read analysis (see Fig. 2b, step 2). Reads with greater than 20 different high-quality candidate alignments are also removed from discordant read analysis since the true location of the read is ambiguous. The remaining discordant reads are clustered based on their orientation and pair-mate locations. Regardless of the results of the discordant read realignment strategy, the sequences of all candidate discordant reads are used in the local assemblies.

### Pair-mate lookup and assembly window merging

To improve the power for detecting large rearrangements and SVs with breakpoints separated by more than the size of the local assembly window, candidate partner loci are identified from the discordant read clusters used to indicate additional genomic loci to extract reads from prior to assembly. This also provides information about the mapping quality of the pair-mates of discordant reads, which is not typically stored in the alignment records of individual reads. To reduce the number of lookups of candidate partner loci in the BAM, SvABA uses a default threshold of six discordant reads to trigger a candidate lookup, or three reads from the case BAM when run in case-control mode. The number of reads required to trigger a partner loci lookup can be modified with the-L flag.

### Sequence assembly with SGA

SvABA performs sequence assembly using a custom version of the String Graph Assembly (SGA) pipeline (Simpson and Durbin 2012) that performs FM-indexing, string graph construction and graph traversal entirely in memory with no intermediate files. The SGA implementation in SvABA deviates from the default settings for whole-genome *de novo* assemblies in order to improve the sensitivity for building variant-spanning contigs. The default in SvABA is to require a minimum overlap of 60% of the read length when evaluating overlaps between reads, to not trim any terminal branches of the assembly graphs, and to exclude contigs with length less than 130% of the read length.

The command line version of SGA produces a text file (ASQG file) representation of the overlap graph, which can be used to visualize and debug assemblies. To allow for rapid debugging and visualization of the local assemblies, SvABA optionally emits the ASQG files for each local assembly window using the -write-asqg option. Because this will produce an ASQG file for each assembly window, the recommended use is to produce ASQG files only for debugging targeted assemblies.

### Contig alignment and candidate variant generation

Following assembly, SvABA aligns the assembly contigs to the human reference using BWA-MEM(Li 2013) and searches for evidence of variant-supporting alignments. To improve the probability that a contig supporting a true variant will correctly align with a gapped or multi-part alignment, SvABA uses the following BWA-MEM options: gap-opening penalty of 32, gap extension penalty of 1, mismatch penalty of 18, match score of 2, Smith-Waterman bandwith of 1000, reseed trigger of 1.5, and 3’ and 5’ clipping penalties of 5. The effect of these parameters is to allow for alignment with large gaps (as found in large indels and SVs), and to minimize excessive single nucleotide mismatches that could create false positive alignments. Regardless of whether the contig alignments support a candidate variant, the alignment records of all of the contigs are emitted directly to a BAM file.

The most conservative alignment for a contig is the one that aligns to within the local window from where the reads were extracted and with no candidate variant. To explicitly check for this possibility, the reference sequence from the local assembly window is extracted and indexed with BWA-MEM. Contigs are aligned to this local reference and excluded from further consideration if they have a high quality non-variant local alignment with fewer than 30 non-aligned bases and no alignment gaps.

Candidate indels are extracted from contigs that align to the reference with a gapped alignment, and candidate SVs are extracted from contigs with mutli-part alignments (see Fig. 2b). High-quality secondary alignments, where a sequence fragment has multiple possible alignments for the same bases, are retained if they have an alignment score (AS score) of greater then 90% of the maximum AS score, up to a maximum of 50 alignments. Although these alignments may support true variants, they are inherently ambiguous and likely overlap repetitive elements that are present at more than one copy in the reference genome. SvABA handles these contigs by reporting all of the candidate variants, one for each of the possible secondary alignments, in an unfiltered VCF (**Supplementary Fig. 2**). These candidate rearrangements can then be disambiguated using copy-number data or other genome-wide analyses to select the most likely variant from the set of candidates.

### Realignment of sequence reads to assembly contigs

To obtain the read support for a candidate variant, within each assembly window all of the reads are aligned to both the contigs and the local reference sequence using BWA-MEM. To be considered a match to a contig, a read must have an AS score of >90% of the length of the match, and have a higher alignment score to the contig than the reference. Clipped read-to-contig alignments are also considered, but only the matched portion is used to indicate read support. Alignment positions and CIGAR strings of the read-to-contig alignments are stored as a tag within with the reads and optionally emitted to a BAM file.

Read-to-contig alignments that span a candidate indel or breakpoint are used to obtain the variant read count. Reads that have an alignment of eight bases to the left and right of a variant site are considered a variant-supporting read. For variants that overlap simple repeats (e.g. homopolymer repeats), this minimum read-to-contig coverage is extended by the length of the repeat. To facilitate rapid review of the evidence for a given contig and variant, the read-to-contigs alignments and contig-to-genome emitted as ASCII plots in the *.alignments.txt.gz file (see Fig. 5b).

### Rearrangement and breakpoint annotation

SvABA annotates indels and SVs with breakpoint microhomology, the sequences of breakpoint insertions, and whether the contig contains evidence for templated-sequence insertions (TSI). Microhomology bases are obtained from overlapping BWA alignments on the contig (see Fig. 5b, second breakpoint). Conversely, breakpoint insertion bases are called when there was a gap between the two aligned fragments (see Fig. 5b, first breakpoint). Rearrangements containing three or more alignments to the reference are annotated with a TSI field in the VCF, and represent TSI rearrangements. To be considered a true TSI rearrangement, both the leftmost and rightmost alignments in the contig coordinates must have a minimum BWA mapping quality of 30 and be supported by at least 4 breakpoint-spanning reads.

### Indel variant scoring and filtering

Candidate indels are initially heuristically filtered to exclude variants from contigs with poor BWA mapping quality (< 10), from contig fragments with multiple ambiguous matches, and from contigs with highly uneven coverage of supporting reads (< 80% of contig covered by a high-quality read-to-contig alignment). Indels with an allelic fraction of less than 0.05 are also removed from the final callsets. All candidate indels failing a filter are output in the unfiltered VCF files.

For short read sequencing by synthesis, the likelihood that a read contains an artificial indel is largely determined by the number of repeats at a site, with large homopolymer stretches being most likely to contain false indels (Ross et al. 2013). To obtain an estimate for the probability that a variant-supporting read is an artifact, SvABA measures the number of repeats in the reference genome immediately to the left and right of indel sites. Repeats are calculated by iteratively moving along the reference genome away from the indel until the repeat pattern is broken, for repeat units up to 5 bp. The repetitive sequences are reported in the VCF files. The total length of the repeat is then converted to an error rate estimate *e* provided in Ross *et al*(Ross et al. 2013).

The remaining indels are scored by calculating the log-of-odds (LOD) that a variant has a non-zero allelic fraction *f* versus homozygous reference (*f =* 0)

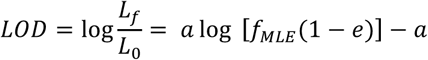

where *f*_*MLE*_ is the maximum likelihood estimate for *f* obtained from the number of variant supporting reads *a* divided by the total number of reads *k*. The default LOD cutoff is 8, or 6 if the variant is present in the dbSNP database.

SvABA will classify an event as germline or somatic if both case (tumor) and control (paired-normal) BAM files are supplied. This functionality can also be used to call *de novo* variants in trios (mother, father, proband child) or quads. Any number of BAM files can be supplied, and variants will be genotyped for each input sample. To determine if an indel is somatic, we follow a similar approach to the calculations performed by MuTect (Cibulskis et al. 2013). For each candidate somatic indel, the LOD that the indel is homozygous reference in the paired-normal (*f* = 0) rather than heterozygous (*f* = 0.5), is calculated from variant and reference read counts as above. The default LOD cutoff for somatic classification is 6.0, or 10.0 if the variant is present at a dbSNP site (and thus more likely to be a germline variant).

### Genotype calculations

For each variant and input sample, SvABA estimates the genotype log-likelihoods as described in equation 2 of Li (Li 2011):

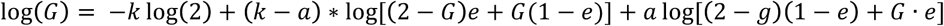

where *G* is the genotype (2 for homozygous reference, 0 for homozygous variant, 1 for heterozygous), *k* is the total number of reads at the variant site, *a* is the number of alternate (variant-supporting) reads and *e* is the estimated error rate at the loci as described above.

### Variant calling from long-sequences using SVlib

The input to SVlib is a BAM file of long-sequence alignments, sorted by the sequence names. SVlib iterates through the BAM and stores any sequence with either a gapped alignment or with multiple alignments (candidate SV) for further processing. The CIGAR strings of the alignments are parsed to extract candidate indel locations in both contig and reference coordinates, and the multipart alignments are parsed to provide candidate SV breakpoints in long-sequence and reference coordinates.

SVlib can optionally genotype the variants by aligning short read sequences to the long-sequences and obtaining the number variant-supporting reads. For each candidate variant, SVlib extracts the reads from the regions covered by the alignment of the long-sequence to the reference. The reads are then aligned to the long-sequences with BWA-MEM, and reads with alignments that span the breakpoints are counted as variant-supporting reads. The genotype calculations and somatic versus germline classifications are performed as described above.

### SV and indel detection in Pacific Biosciences sequences

Whole-genome assemblies of PacBio reads from Pendleton *et al* (Pendleton et al. 2015) were downloaded from Sequence Read Archive (SRX627421 and SRX638310). Contigs were fragmented into 2 kb sequences and aligned to hg19 reference with BWA-MEM 0.7.12 using the following parameters:-Y -A2 -E1 -O32 -B19. SVlib was then used to parse the gapped and multiple-alignment contigs as described above. Variant read support and genotyping was performed with SVlib by realigning the 151 base reads to the PacBio contigs.

### Comparison of SV and indel calls

To reduce confounding effects of differing conventions and detection strategies implemented by the various SV and indel detection tools, we allowed some difference between breakpoint locations for different callers when comparing overlaps between callsets. For short events, we considered two events as overlapping if their first breakpoints were within 10 bp of each other. For SVs larger than 50 bp, we allow up to 100 bp of difference between the variant sites, with the added requirement that both the left and right breakpoints of large SVs (> 300 bp) overlap between the callsets. Because the spacing between large SVs in the human genome is typically on the order of 10-100s of kb, the rate of spurious overlaps due to allowing a small amount of breakpoint ambiguity is insignificant. For all methods, calls from regions with very low sequence mappability were excluded using the regions defined in: https://data.broadinstitute.org/snowman/svaba_exclusions.bed.

### In silico tumor genome

SVlib generated an *in silico* tumor genome by creating rearranging the reference genome (hg19) and generating 2,000 bp contigs spanning the rearrangements. Virtual reads were generated from the breakpoint contig FASTA files using the ART read simulator (Huang et al. 2012) and the standard Illumina error profile. We simulated 101 base pair read pairs at 10x coverage across the breakpoint contigs. To simulate a heterozygous, impure tumor genome, we combined the simulated reads with real sequence reads from the HCC1143 lymphoblastoid normal cell line (HCC1143_BL). Read-pairs from HCC1143_BL were randomly selected to achieve an average sequence coverage of 28.8. To simulate the matched normal, we further randomly selected read-pairs from HCC1143_BL to obtain a subsampled BAM at a mean coverage of 57.7, with no read-pairs in common with the 28.8x BAM. All reads were aligned with BWA-MEM using default parameters with the SpeedSeq pipeline (Chiang et al. 2015).

### Identifying sites of viral integration

The Refseq1.1 database was downloaded from ftp://ftp.ncbi.nlm.nih.gov/refseq/release/viral/ and indexed with BWA. Assembled contigs from each local assembly window were mapped to both the human reference and viral database. To enrich for only junction-supporting contigs, contigs mapping entirely to the viral database were excluded. Only contigs with ≥ 50 bases unmapped to the human reference and ≥ 35 bases mapping to the viral reference were considered as candidates for supporting a viral insertion junction. The contigs were parsed to obtain the genomic location of the integration site, and scored to obtain read support using the same procedure as for non-viral SVs. The viral integration calls are then emitted directly in the VCF files. A BAM file containing all of the viral alignments can be optionally emitted.

## Data and software availability

SvABA is freely available under the GPLv3 license at https://github.com/walaj/svaba, and contains extensive documentation and several use-case examples. SVlib is available under the GPLv3 license at https://github.com/walaj/svlib.

The 151 base paired-end Illumina sequencing data on NA12878, the HCC1143 and HCC1143_BL 101 base paired-end reads and 250 base PCR-free paired-end reads, and Discovar *denovo*assemblies are publically available at https://data.broadinstitute.org/snowman/sequences (Sequence Read Archive submission pending). The variant calls for SVlib, SvABA, DELLY, LUMPY and Pindel on NA12878 are availableat: https://data.broadinstitute.org/snowman/Submission/NA12878.

The simulated tumor BAM files, simulated tumor genomes, truth set of variant calls, and SvABA, DELLY, LUMPY, FreeBayes, Pindel and Strelka calls from the simulated data are available at. https://data.broadinstitute.org/snowman/Submission/Simulations/SimData/. The TCGA whole-genomes were obtained from the NIH GDC portal (https://gdc-portal.nci.nih.gov/). The TCGA external IDs of the submitted samples are listed in **Supplementary Table 5**.

## Acknowledgements

The National Institutes of Health (T32 HG002295/HG/ NHGRI, U54CA143798, and R01CA188228), DFCI-Novartis Drug Discovery Program, Voices Against Brain Cancer, Pediatric Low-Grade Astrocytoma Foundation, the Broad Institute, and the Cure Starts Now Foundation provided financial support. We would like to thank Heng Li for helpful comments and as the primary developer of BWA-MEM, and Jared Simpson as the developer of SGA.

### Author contributions

J.A.W., M.I., C.Z., M.M., R.B. wrote the paper. J.A.W. developed SvABA and performed analyses. P.B. and R. O. performed the HCC1143 validation experiments. N.G., C.S., Y.L., J.W., P.C., S.S. contributed methodological improvements. X.Y. performed the TCGA variant calling, T.S. and C.N. performed the HCC1143 data generation and Discovar *de novo* assemblies. M.I., C.Z., and R.B. supervised the research.

### Disclosure declaration

The authors declare that there is no conflict of interest.

